# A versatile platform for single fluorescent protein-based fluorescence lifetime biosensors

**DOI:** 10.1101/2024.06.29.601303

**Authors:** Chongxia Zhong, Satoshi Arai, Yasushi Okada

## Abstract

Single fluorescent protein (FP)-based FLIM (fluorescence lifetime imaging) biosensors are potent tools for quantitatively imaging intracellular processes with high spatial and temporal resolution. They only require a single wavelength for detection, which facilitates multi-color imaging. However, the development of single FP-based FLIM biosensors has been limited by the absence of a general design framework and the complexity of the screening process. In this study, we engineered FLIM biosensors capable of detecting ATP (adenosine triphosphate), cAMP (cyclic adenosine monophosphate), citrate, and glucose by inserting each sensing domain into the mTurquoise2 fluorescent protein between Tyr-145 and Phe-146 using peptide linkers. Through efficient linker screening, we successfully developed FLIM biosensors exhibiting an effective dynamic range from 0.5 to 1.0 ns upon analyte binding. This demonstrates that the qmTQ2-ATP-0 backbone is a universal platform for developing mTQ2-based biosensors. As a proof-of-concept, we demonstrated the capabilities of these FLIM biosensors in monitoring the intracellular dynamics of ATP and cAMP alongside dual-color imaging. Therefore, our work presents an accessible methodology for establishing a single FP-based FLIM biosensor platform for quantitative imaging.

## Introduction

Fluorescent protein (FP)-based biosensors play a crucial role in cell biology by offering dynamic insights into the localization and quantity of signaling molecules and metabolites. Currently, fluorescent intensity-based biosensors are widely used by the two designs: single FP-based intensiometric biosensors and two-FP-based FRET biosensors. The single FP-based intensiometric biosensors report their target analytes based on changes in fluorescence intensity (Greenwald et al., 2018; Nasu et al., 2021). However, intensiometric biosensors are susceptible to various experimental factors affecting the fluorescent signal intensity, such as photobleaching, probe concentration, excitation intensity, and sample thickness, compromising measurement accuracy. FRET-based biosensors use the emission ratio of two FPs to avoid these issues and report the analyte more quantitatively (Yellen & Mongeon, 2015). Cyan FP (CFP) and yellow FP (YFP) pairs have been versatile platforms for FRET-based biosensors for various analytes or reactions (Zhang et al., 2001) (Klarenbeek et al., 2015) (Terai et al., 2019). However, the FRET-based biosensors use two FPs for one biosensor, which limits the multiplex measurement with multiple biosensors for different analytes. Moreover, the differences in the maturation and degradation kinetics of the two FPs can bias the measurements (Yaginuma et al., 2014).

Fluorescence lifetime imaging microscopy (FLIM) provides a more robust quantitative imaging approach, as fluorescence lifetime—an intrinsic property of the fluorophore—is not affected by the aforementioned experimental factors (Datta et al., 2020). Genetically encoded FLIM biosensors fall into two categories: FRET-FLIM biosensors and single FP-based FLIM biosensors. FRET-FLIM biosensors utilize changes in fluorescence lifetime caused by energy transfer between a donor and an acceptor fluorophore to detect and quantify analytes (Kukk et al., 2022). However, its design to use two FP pairs poses similar limitations with the intensity-based FRET biosensors mentioned above: the potential bias by the differences in maturation and degradation rates between the two FPs and the limitations in multiplex measurement.

Single FP-based FLIM biosensors can overcome these drawbacks of the FRET-FLIM biosensors (Vu & Arai, 2023). Several single FP-based FLIM biosensors have been engineered so far, such as FLIM biosensors for sensing the NADH/NAD^+^ ratio (Hung et al., 2011), ATP (Arai et al., 2023), glucose (Diaz-Garcia et al., 2019), Ca^2+^(van der Linden et al., 2021) and lactate (Koveal et al., 2022). However, the number of single FP-based FLIM biosensors is limited compared to intensity-based biosensors. This would be partly due to the lack of a design framework and the technical requirements for screening, such as the automated FLIM microscope.

This study aims to expand the FLIM biosensor toolbox by developing single FP-based FLIM biosensors through an easy-to-implement screening approach combined with a versatile platform. We chose a CFP, mTQ2, as the scaffold for constructing FLIM biosensors to sense intracellular signaling molecules. mTQ2 boasts a long mono-exponential fluorescence lifetime (4.0 ns) and a high quantum yield (0.93) (Goedhart et al., 2012). The long mono-exponential fluorescence lifetime of mTQ2 was achieved by ensuring the planarity of the chromophore and restricting its rotational freedom through strong van der Waals interactions with surrounding amino acids (Goedhart et al., 2012). Therefore, we expected perturbations to the beta-can amino acid residues surrounding the chromophore would increase the non-radiative relaxation, decreasing the lifetime and the quantum yield. Additionally, the fluorescence of mTurquoise2 remains stable across a wide pH range due to its low pKa of 3.1, suggesting that mTQ2-based fluorescent biosensors may show high stability against physiological pH fluctuations. Reported mTQ2-based FLIM biosensors, such as Tq-Ca-FLITS and Lilac, alter both intensity and lifetime by analytes (Koveal et al., 2022; van der Linden et al., 2021). This would reflect that mTQ2 has a minimal non-radiative decay (long lifetime and high quantum yield). For other FPs, changes in fluorescence intensity and lifetime may not necessarily coincide, as their non-radiative decay rates might be higher and more variable. Lifetime changes do not correlate well with the intensity changes in other fluorescent biosensors, such as the GFP-based GCaMP6 and jGCaMP7 (Dana et al., 2019) and the RFP-based R-GECO1 (Zhao et al., 2011).

In this paper, we propose mTQ2 as a versatile platform for single FP-based FLIM biosensors suitable for a fluorescence intensity-based screening approach. As a proof-of-concept, we first examined the properties of an mTQ2-based FLIM biosensor for ATP. Subsequently, we validated our biosensor design platform and screening strategy by developing FLIM biosensors for sensing cAMP, citrate, and glucose.

## Materials and Methods

### Chemicals and Reagents

ATP, adenosine diphosphate (ADP), cAMP, disodium citrate sesquihydrate, and histamine dihydrochloride were purchased from Tokyo Chemical Industry Co., Ltd. (Tokyo, Japan). Forskolin, D (+)-glucose, sodium L (-)-malate, DMEM (Dulbecco’s modified Eagle’s medium), L-lactate, galactose, and penicillin-streptomycin were purchased from WAKO Pure Chemical Industries (Osaka, Japan). Cyclic guanosine monophosphate (cGMP) was purchased from Merck Millipore (Darmstadt, Germany). (-) isoproterenol hydrochloride was purchased from Sigma Aldrich (Missouri, USA). 2-Deoxyglucose, guanosine triphosphate (GTP), deoxyadenosine triphosphate (dATP), adenosine monophosphate (AMP), sodium pyruvate, D-Glucose-1-phosphate, D-Glucose-6-phosphate, 2-Ketoglutaric acid and D-fructose were purchased from Nacalai Tesque (Kyoto, Japan). All oligonucleotides for primers, TrypLE Express, phosphate-buffered saline (PBS), Hank’s balanced salt solution (HBSS) (+), and fetal bovine serum (FBS) were purchased from ThermoFisher Scientific (Massachusetts, USA). KODone PCR master mix and In-Fusion Snap Assembly Master Mix were purchased from Takara Bio Inc. (Shiga, Japan). Restriction enzymes were purchased from New England Biolabs (Massachusetts, USA).

### Plasmid construction

The cDNA fragment for mTurqoise2 (Addgene plasmid #54843) was modified by overlapping extension PCR (polymerase chain reaction) to include KpnI and EcoRI restriction sites between Tyr-145 and Phe-146 of mTQ2. The resultant DNA was cloned into the XhoI/HindIII site of the pRSET-A vector. The resulting construct is named mTQ2-pRSET-A plasmid. For the qmTQ2-ATP biosensor cDNA construction, the epsilon subunit of the bacterial F_o_F_1_-ATP synthase cDNA (Addgene plasmid #113906) was inserted into mTQ2-pRSET-A plasmid at Tyr-145 (between the KpnI and EcoRI restriction sites) through various peptide linkers generated by PCR, using In-Fusion Snap Assembly Master Mix. A similar strategy was adopted for constructing the plasmids of the qmTQ2-cAMP, qmTQ2-citrate, and qmTQ2-glucose biosensors. The sensing domain for cAMP comprised residues of 199–358 of mouse Epac1 (Addgene plasmid #73938). The sensing domain for citrate comprised residues 4–133 of the CitAP domain from *Klebsiella pneumoniae* CitA protein (Addgene plasmid #134301). The sensing domain for glucose comprised residues 24–328 of the bacterial D-galactose-binding periplasmic protein (MglB) (Addgene plasmid #163115). Site saturation mutagenesis was carried out according to a previous protocol (Liu & Naismith, 2008) (Steffens & Williams, 2007). Briefly, sense and antisense primers were created for the targeted amino acid, and the codon was replaced with NNK and MNN, respectively. For mammalian expression, the Kozak sequence was added to the cDNA of qmTQ2-ATP and qmTQ2-cAMP by PCR, and the resulting PCR fragments were cloned into pcDNA3.1(-) vectors using In-Fusion Snap Assembly Master Mix. Kozak sequence was added to RCaMP1h by PCR using RCaMP1h cDNA from pRSET-RCaMP1h plasmid (Addgene plasmid #42874) as the template, and the result PCR product was inserted into the mammalian expression pN1 vector.

### Screening

The mutants for fluorescence biosensor candidates containing plasmids were transformed into *Escherichia coli* JM109 (DE3) competent cells and cultured by shaking at 200 rpm for 3 days at 20°C. Subsequently, the bacterial suspension was harvested by centrifugation at 4440 g for 10 min at 4°C, and the pellet was resuspended in 20 mM HEPES buffer. This was followed by sonication for 30 s (amplitude 60%) using an ultrasonic disruptor (UD-100, TOMY Digital Biology Co., LTD, Tokyo, Japan). The supernatant obtained after centrifugation (17750 g, 20 min) was transferred to a new Eppendorf tube. The fluorescence property of each candidate without or with 10 mM ATP (final concentration) was examined using a microplate fluorescence reader (VarioSkan Lux, ThermoFisher Scientific). For the emission spectra, the excitation wavelength was set to 450 nm, and the fluorescence emission range was from 470 nm to 600 nm at a 1 nm step size. For the excitation spectra, the emission wavelength was set at 520 nm, with the range of excitation from 350 nm to 500 nm at 1 nm step size. The fluorescence lifetime response of the mutants that showed changes in fluorescence intensity upon ATP addition was measured under a Leica TCS-SP8 FALCON confocal FLIM microscope (Leica Microsystems, Wetzlar, Germany). Samples were excited with a 440 nm pulsed diode laser (PicoQuant, Berlin, Germany), and photon arrival times were recorded with a HyD detector, with emission ranging from 470 nm to 530 nm. The same approach was adopted in creating qmTQ2-cAMP, qmTQ2-citrate, and qmTQ2-glucose biosensors. The buffer for resuspending the bacterial pellets is PBS for both qmTQ2-cAMP biosensor and qmTQ2-glucose biosensor, and TBS (Tris-buffered Saline) for qmTQ2-citrate biosensor.

### Fluorescence protein expression, purification, and In vitro characterization

For protein expression, qmTQ2-ATP, qmTQ2-cAMP, qmTQ2-citrate, and qmTQ2-glucose in the pRSET-A vector were transformed into JM109 (DE3) competent cells, respectively. The cells were then cultured by shaking at 200 rpm for 3 days at 20°C and harvested by centrifugation at 4440 g for 10 minutes. The harvested cells were resuspended in PBS, the cell suspensions were sonicated, centrifuged, and supernatants were collected. The supernatants containing the His-tagged fluorescent biosensors were purified with TALON Metal Affinity Resins (Takara Bio USA, Inc., California, USA) according to the manufacturer’s procedure. Proteins were eluted with a buffer containing 50 mM Tris (pH 7.4), 300 mM NaCl, and 200 mM imidazole. Subsequently, the proteins were subjected to buffer exchange to 20 mM HEPES (for qmTQ2-ATP), PBS (for qmTQ2-cAMP and qmTQ2-glucose biosensors), and TBS (for qmTQ2-citrate biosensor) using the PD-10 gel filtration column (GE Healthcare, Illinois, USA) to remove imidazole. The absorption spectra of the purified fluorescent biosensors were measured between 350 nm and 600 nm using a UV spectrophotometer (UV-1800, Shimadzu, Kyoto, Japan) with the buffer as a reference. The emission and excitation spectra were recorded using a fluorescence spectrophotometer (FP-8500, JASCO, Tokyo, Japan). For the excitation spectra, the emission wavelength was set at 520 nm, and the range of excitation from 350 nm to 500 nm. For the emission spectra, the excitation wavelength was 450 nm, and the range of fluorescence emission was from 470 nm to 600 nm.

### Quantum yield and molar absorption coefficient measurements

The quantum yields (QY) of the purified qmTQ2-ATP, qmTQ2-cAMP, qmTQ2-citrate, and qmTQ2-glucose proteins were measured with the absolute PL quantum yield spectrometer (C9920-02, Hamamatsu Photonics K.K., Shizuoka, Japan) with the excitation wavelength set at 450 nm.

The molar absorption coefficient (ε) was determined through the alkaline denaturation method. To achieve this, the concentration of the purified qmTQ2-ATP, qmTQ2-cAMP, qmTQ2-citrate, and qmTQ2-glucose proteins in the presence or absence of the analytes was adjusted to attain a native optical density (OD) of at least 0.2 after 1:1 dilution with buffer. The absorbance spectra were measured before and more than 5 minutes after adding 2 M NaOH (at a 1:1 ratio) across the range of 350–650 nm with a 1 nm step size, using the corresponding buffer as a reference. The concentration of the unfolded protein was determined using the Beer-Lambert law, assuming the molar absorption coefficient at 462 nm of 46000 M^−1^cm^−1^ for the free cyan chromophore (Lelimousin et al., 2009). Subsequently, utilizing the concentration derived from the NaOH-denatured sample, the peak extinction coefficient value for the native samples, in the presence or absence of the analytes at 450 nm, was determined.

### Cell culture and Transfection

HeLa cells (ATCC CCL-2) and COS7 cells (ATCC CRL-1651) were cultured in Dulbecco’s modified Eagle’s medium (DMEM) supplemented with 10% fetal bovine serum, 100 U/ml penicillin and 100 μg/mL streptomycin at 37°C under a humidified atmosphere containing 5% CO_2_. HeLa cells were cultured in low-glucose DMEM. COS7 cells were cultured in high-glucose DMEM. For imaging experiments, the cells were dissociated with TrypLE and then seeded onto 35 mm glass-bottom dishes (P35G-1.5-14-C, MatTek Life Sciences, Massachusetts, USA) coated with collagen I. Once the cells reached 70% confluency, they were transfected with plasmid in the pcDNA3.1(-) vector (0.2 μg plasmid DNA per dish) using TransFectin Lipid Reagent (#170350, Bio-Rad Laboratories Inc., California, USA) according to the manufacturer’s instructions. The medium was exchanged 4 hours after transfection, and the cells were cultured at 37°C for 24 hours before imaging.

### Fluorescence Lifetime Imaging

FLIM experiments were performed at room temperature using a Leica TCS-SP8 FALCON confocal FLIM microscope with LAS-X version 3.5 software and a 93× objective lens (HC PL APO CS2 93×1.30 LYC). Samples were excited with a 440 nm pulsed diode laser, and photon arrival times were recorded using the HyD detector, with the emission ranging from 470 nm to 530 nm. Images were acquired every 5 seconds for 20 minutes in the forskolin and isoproterenol treatment experiments. Images were acquired every 10 seconds for 15 minutes for the qmTQ2-cAMP and RCaMP1h dual-color imaging experiments. The initial treatment commenced two minutes after starting image acquisition.

Fluorescence lifetimes were calculated using a standardized “tau8” value proposed by the Yellen group (Diaz-Garcia et al., 2019). The fluorescence lifetime decay histogram was fitted with a two-exponential function, and the photon arrival time was fixed to 0–8 ns from the start of the pulse. Subsequently, the intensity-weighted average lifetime (Tau int) was reported.

### Data Analysis

Time-lapse results of fluorescence lifetime in cells were analyzed using ImageJ. GraphPad Prism 10 was used to plot the generated data and perform statistical analysis.

## Results

### Construction, characterization, and validation of the FLIM biosensor for ATP

ATP provides energy to drive many essential biological processes in living organisms. ATP biosensors are valuable tools for evaluating the spatiotemporal dynamics of ATP within a single cell. We decided to begin with ATP FLIM biosensor construction because our team has previously established methods for engineering intensiometric ATP biosensors (Arai et al., 2018), and a YFP-based FLIM biosensor for ATP detection was reported recently by our group (Arai et al., 2023). An mTQ2-based ATP FLIM biosensor would expand the color variants and offer potential for in vivo imaging using two-photon microscopy. We constructed the ATP FLIM biosensor by inserting the epsilon subunit of *B. subtilis* F_o_F_1_ ATP synthase into mTQ2 protein between Tyr-145 and Phe-146 and designated it as qmTQ2-ATP-0 (quantitative mTurquoise2-based ATP biosensor version 0) (Figure S1A). The epsilon subunit of F_o_F_1_ ATP synthase specifically binds to ATP without hydrolyzing it and undergoes a significant conformational change upon ATP binding (Arai et al., 2018; Imamura et al., 2009). The insertion site was determined according to previous studies (Baird et al., 1999; Matsuda et al., 2017) to ensure that the sensing domain would be in close proximity to the chromophore of mTQ2 upon folding into its tertiary structure. We expected that the ATP-dependent conformational change in the epsilon subunit would influence the chromophore’s environment, resulting in alterations in its fluorescence properties, including fluorescence lifetime.

The fluorescence intensity or lifetime of bacterial lysate of qmTQ2-ATP-0 did not change in the presence of 10 mM ATP. Previous studies indicated that modifying the linker length and/or the type of amino acids in the linker sequence between the sensing domain and the fluorescent protein could enhance the dynamic range of fluorescent indicators (Odaka et al., 2014). Therefore, we performed linker screening on the qmTQ2-ATP-0 construct. The amino acid sequence (ALKKEL) from the NZ leucine zipper motif (Ghosh et al., 2000) was adopted as the basis of the N-terminus linker (linker 1) and C-terminus linker (linker 2). We anticipated that the strong alpha-helix structure of the NZ leucine zipper would facilitate a more efficient transfer of the conformational change of the epsilon subunit to the chromophore, thereby improving the dynamic range of the ATP FLIM biosensor (Arai et al., 2018). Linker libraries were created with 0 to 6 amino acid insertions in both linkers (Figure S1A). Screening the bacterial lysates from the linker library for ΔF/F using a fluorescence plate reader led to identifying a mutant, which showed a 5% fluorescence intensity change in response to 10 mM ATP (Figure S1C). This mutant, named qmTQ2-ATP-0.1, contains a 5-amino-acid insertion (ALKKE) in linker 2 (Figure S1B).

Previous studies reported that Tyr-145 in green fluorescent protein (GFP) plays an important role in enhancing the performance of fluorescent biosensors since it is sterically adjacent to the chromophore (Baird et al., 1999). To expand the dynamic range of the qmTQ2-ATP-0.1 mutant, further optimization was pursued through site saturation mutagenesis targeting Tyr-145. Initially, we screened the bacterial lysates for ΔF/F using the fluorescence plate reader. Subsequently, we measured the fluorescence lifetime change (Δτ) of the bacterial lysates from the mutants showing significant ΔF/F. The screening yielded a mutant displaying a 19% reduction in fluorescence intensity and a 0.6 ns decrease in fluorescence lifetime in response to 10 mM ATP. This mutant, designated as qmTQ2-ATP-0.2, featured the Y145K substitution (Figures S1D–S1G).

Subsequent site-saturation mutagenesis targeting the glycine residue of linker 1 resulted in the identification of qmTQ2-ATP-0.3, which showed a 25% decrease in fluorescence intensity and 0.9 ns reduction in fluorescence lifetime upon exposure to 10 mM ATP (Figures S1H–S1K). In qmTQ2-ATP-0.3 the glycine of linker 1 was mutated to proline alongside the Y145K substitution. Using qmTQ2-ATP-0.3 as the template, a third round of site-saturation mutagenesis targeting the threonine residue of linker 1 was conducted, and a mutant with a 39% fluorescence intensity decrease and 1.1 ns decrease in fluorescence lifetime in the presence of 10 mM ATP was obtained. This mutant, featuring substitutions at glycine (to proline) and threonine (to valine) of linker 1, in addition to the Y145K mutation, was named qmTQ2-ATP biosensor (Figure 1A).

**Figure 1.**
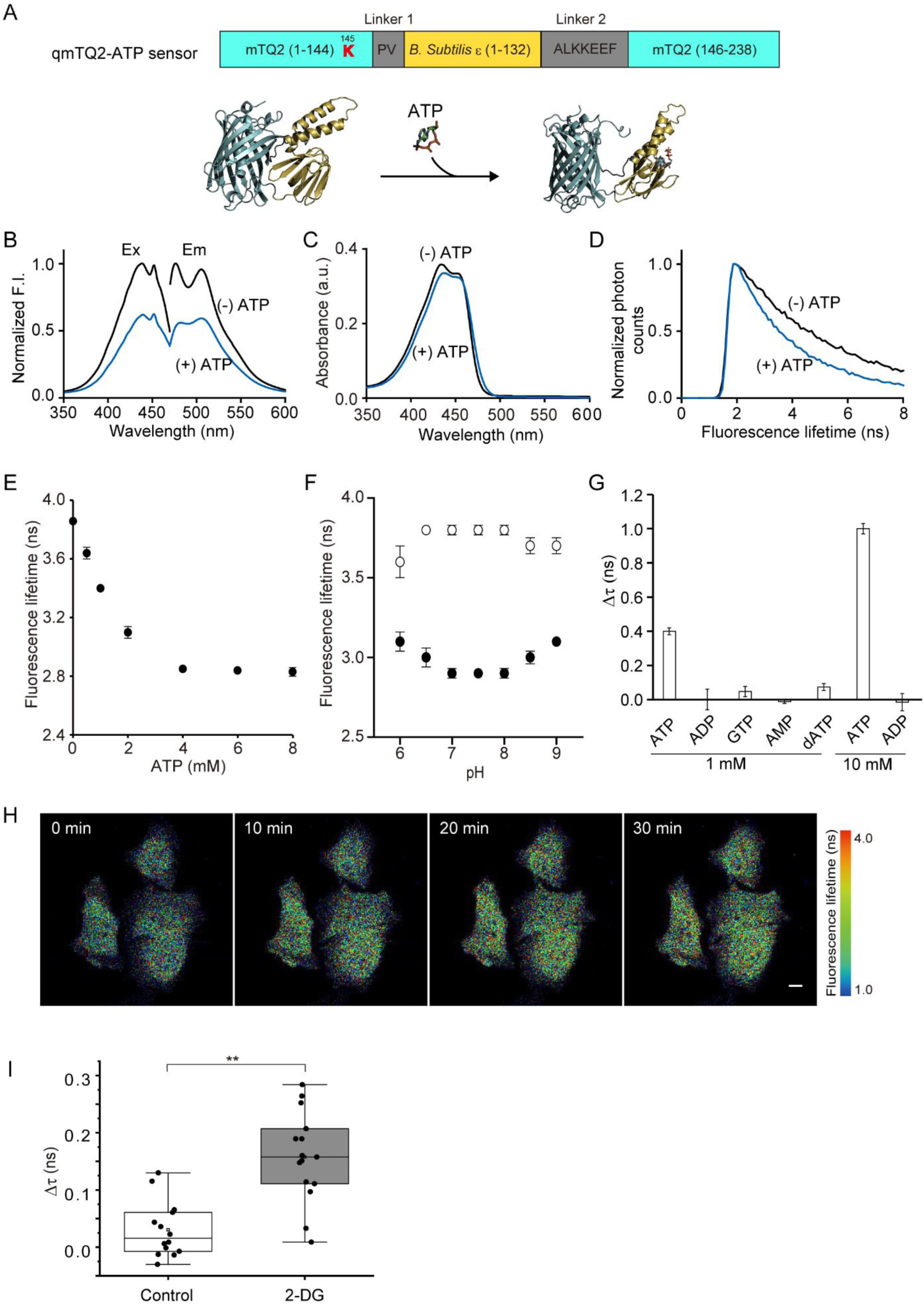
Construction, characterization, and validation of qmTQ2-ATP biosensor. (A) Schematic representation of the structural domain and three-dimensional (3D) structural models of qmTQ2-ATP biosensor. 3D structure models were generated with Alphafold2 (Jumper et al., 2021). (B–D) The emission and excitation spectra (B), the absorption spectra (C) and the fluorescence decay curve (D) of qmTQ2-ATP biosensor in the presence (blue) and absence (black) of 10 mM ATP. (E) The dose-response curve of qmTQ2-ATP biosensor to ATP in solution. The data represents means ± SD (n = 5). (F) Effect of pH on the fluorescence lifetime of qmTQ2-ATP biosensor in the presence (filled circle) and absence (open circle) of 10 mM ATP. The data represents means ± SD (n = 5). (G) Specificity of qmTQ2-ATP biosensor to ATP and other nucleotides. Δτ represents the lifetime changes with the presence and absence of nucleotides. The data represents means ± SD (n = 5). (H) Sequential pseudo-color images of HeLa cells expressing qmTQ2-ATP biosensor in response to 20 mM 2-DG. Fluorescence lifetime (τ) with pseudo color, scale bar: 10 μm. (I) Box-whisker plot comparing Δτ in HeLa cells between the untreated control group and the group treated with 2-DG for 30 minutes. Double asterisks indicate p<0.05 by Student’s t-test.

The results of in vitro characterization with purified protein revealed that both the emission and excitation spectra of the purified qmTQ2-ATP biosensor exhibited a 39% reduction in amplitude upon saturation with ATP, with a fluorescence lifetime change from 3.8 ns (ATP-free state) to 2.7 ns (ATP-bound state). Notably, the absorption spectra of qmTQ2-ATP biosensor remained comparable in the presence or absence of ATP (Figures 1B–1D, Table 1). Furthermore, the quantum yield of the qmTQ2-ATP biosensor decreased from 0.71 to 0.47 upon ATP addition, while the molar extinction coefficient showed only a slight change (Table 1). The fluorescence lifetime of qmTQ2-ATP biosensor altered in response to ATP in a dose-dependent manner, with a dissociation constant (K_d_) estimated at 1.3 ± 0.2 mM (Figures 1E and S2). Additionally, the fluorescence lifetime of the qmTQ2-ATP biosensor remains insensitive to pH changes within the range of 7.0 to 8.0 (Figure 1F), indicating its suitability for quantitative intracellular imaging. Moreover, qmTQ2-ATP showed specificity to ATP compared to other nucleotides (Figure 1G).

**Table 1.** Photophysical characterization of the FLIM sensors developed in this paper.

| Name | Analytes | $\tau$<br>(ns) | $\epsilon$<br>($\times 10^4 \text{ M}^{-1} \text{ cm}^{-1}$) | QY | $k_r$<br>($\times 10^8; \text{ s}^{-1}$) | $k_{nr}$<br>($\times 10^8; \text{ s}^{-1}$) |
| --- | --- | --- | --- | --- | --- | --- |
| qmTQ2-ATP | 10 mM ATP | 2.7 | 2.61 | 0.47 | 1.69 | 1.90 |
|  | 0 mM ATP | 3.8 | 2.69 | 0.71 | 1.83 | 0.76 |
| qmTQ2-cAMP | 1 mM cAMP | 3.6 | 3.18 | 0.65 | 1.80 | 0.98 |
|  | 0 mM cAMP | 3.0 | 3.25 | 0.51 | 1.70 | 1.63 |
| qmTQ2-citrate | 20 mM citrate | 3.0 | 3.20 | 0.45 | 1.50 | 1.83 |
|  | 0 mM citrate | 2.2 | 2.72 | 0.32 | 1.45 | 3.10 |
| qmTQ2-glucose | 10 mM glucose | 2.5 | 3.86 | 0.37 | 1.48 | 2.52 |
|  | 0 mM glucose | 3.0 | 3.92 | 0.52 | 1.73 | 1.60 |

$\tau$ , Fluorescence lifetime; $\epsilon$ , Molar absorption coefficient; QY, Quantum yield.

Next, we validated the performance of the qmTQ2-ATP biosensor by expressing it in HeLa cells. Thirty minutes following the treatment with 20 mM 2-Deoxy-D-glucose (2-DG), a glycolysis inhibitor, the fluorescence lifetime of qmTQ2-ATP increased by about 0.1 ns compared to the control group, indicating that qmTQ2-ATP could detect changes in intracellular ATP levels induced by glycolysis inhibition (Figures 1H and 1I).

The successful development of qmTQ2-ATP biosensors confirms our hypothesis that an mTQ2-based FLIM biosensor can be developed using a fluorescence intensity-based screening approach. Similar to Tq-Ca-FLITS, the changes in fluorescence intensity and fluorescence lifetime of qmTQ2-ATP mutants are also closely correlated (Table S1). Next, we explored the feasibility of using qmTQ2-ATP-0 as a generalizable backbone to engineer a suite of mTQ2-based FLIM biosensors by substituting the ATP sensing domain with the sensing domains for other signaling molecules and metabolites.

### Construction, characterization, and validation of the cAMP FLIM biosensor

cAMP is a widely conserved intracellular second messenger. Its concentration is tightly controlled by the balance of adenylyl cyclases (ACs) and phosphodiesterase (PDE) activities in mammalian cells (Kawata et al., 2022). Jalink’s research group engineered Epac (the exchange protein directly activated by cAMP)-based FRET FLIM biosensors to measure changes in cellular cAMP levels (Ponsioen et al., 2004) (Klarenbeek et al., 2015) (Kukk et al., 2022). To our knowledge, a single FP-based FLIM indicator for cAMP has not been reported.

We engineered an mTQ2-based cAMP FLIM biosensor by inserting the cAMP binding domain of mouse EPAC1 (Odaka et al., 2014) into mTQ2 between Tyr-145 and Phe-146. The resulting construct, qmTQ2-cAMP-0 (quantitative mTurquoise2-based cAMP biosensor version 0) (Figure S3A), showed a 1.4% increase in fluorescence intensity upon adding 1 mM cAMP. Through screening the bacterial lysate of the linker library for ΔF/F_0_, a mutant that showed an 8% decrease in fluorescence intensity in response to 1 mM cAMP was found (Figure S3B). This mutant, designated as qmTQ2-cAMP-0.1, contains a 2 amino-acid (AL) insertion in linker 2 (Figure S3A). Following site saturation mutagenesis aimed at Tyr-145 of qmTQ2-cAMP-0.1 led to the identification of a mutant displaying a 23% elevation in fluorescence intensity and a 0.6 ns increase in fluorescence lifetime upon exposure to 1 mM cAMP. This mutant, carrying the Y145R mutation, was named qmTQ2-cAMP biosensor (Figure 2A).

**Figure 2.**
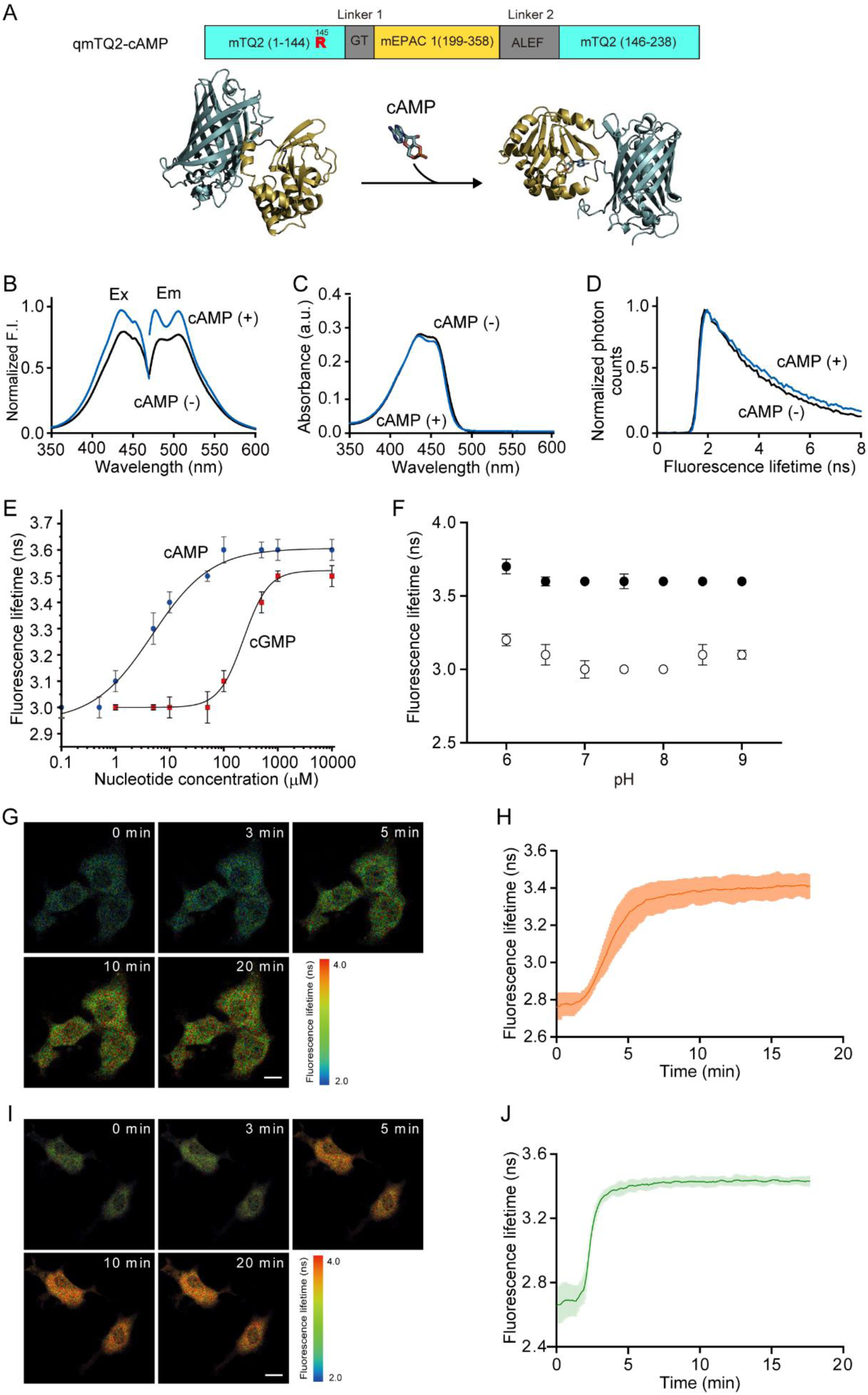
Construction, characterization, and validation of qmTQ2-cAMP biosensor. (A) Schematic representation of the structural domain and 3D structure models of qmTQ2-cAMP biosensor. 3D structure models by Alphafold2. (B–D) The emission and excitation spectra (B), the absorption spectra (C) and the fluorescence decay curve (D) of qmTQ2-cAMP biosensor in the presence (blue) and absence (black) of 1 mM cAMP. (E) Dose-responsive curves of qmTQ2-cAMP biosensor for cAMP (blue circles) and cGMP (red squares). The data represents means ± SD (n = 5). (F) Effect of pH on the fluorescence lifetime of qmTQ2-cAMP biosensor in the presence (closed circles) and absence (open circles) of 1 mM cAMP. The data represents means ± SD (n = 5). (G-H) Representative images (G) and the time course (H) of fluorescence lifetime changes in response to 50 μM forskolin stimulation in qmTQ2-cAMP biosensor expressing COS7 cells (scale bar: 10 μm) (n = 7). (I-J) Representative images (I) and the time course (J) of fluorescence lifetime changes induced by 100 μM isoproterenol application in qmTQ2-cAMP biosensor expressing COS7 cells (n = 7) (scale bar: 10 μm).

The emission and excitation spectra of qmTQ2-cAMP biosensor increased in amplitude by 23% upon the addition of 1 mM cAMP (Figure 2B) accompanied by a fluorescence lifetime change from 3.0 ns (cAMP-free state) to 3.6 ns (cAMP-bound state) (Figure 2D). The alteration in the qmTQ2-cAMP biosensor absorption spectra was small between the presence and absence of cAMP (Figure 2C). The specificity of the qmTQ2-cAMP biosensor for cAMP was examined by performing cAMP and cGMP (cyclic guanosine monophosphate) titration (Figure 2E). The determined K_d_ values for cAMP and cGMP were 5.8 ± 1.2 μM and 228.9 ± 28.3 μM, respectively (Figure S4), indicating high specificity of qmTQ2-cAMP sensor for cAMP and low affinity for cGMP. The range of cAMP concentration detected by this indicator spanned from 1 to 100 μM (Figure 2E), covering both basal and elevated cAMP concentrations reported in mammalian cells (Borner et al., 2011; Kawata et al., 2022). Furthermore, the fluorescence lifetime of qmTQ2-cAMP biosensor remained insensitive to pH changes between 7.0 and 8.0, suggesting its suitability for quantitative intracellular imaging (Figure 2F).

Subsequently, we evaluated the intracellular performance of qmTQ2-cAMP biosensor. In COS7 cells expressing qmTQ2-cAMP, the fluorescence was uniformly distributed in the cytoplasm but excluded from the nucleus (Figure 2G). In Figure 2G, the pseudo-colored images illustrate the FLIM response of the biosensor before and at different time points post-stimulation with 50 μM forskolin. The fluorescence lifetime of qmTQ2-cAMP biosensor increased significantly shortly after forskolin application (Figure 2H), as forskolin activates adenylate cyclase, leading to an elevation in intracellular cAMP concentration. Furthermore, we evaluated qmTQ2-cAMP biosensor in COS7 cells using a β-adrenergic receptors agonist, isoproterenol. The results suggested an increase in the fluorescence lifetime of qmTQ2-cAMP biosensor upon 100 μM isoproterenol stimulation (Figures 2I and 2J), demonstrating its capability to monitor cAMP dynamics in live cells under physiological conditions.

The results indicate that we have identified an insertion site (between Tyr-145 and Phe-146) within mTQ2 that has the capability to transform the interaction signal between the sensing domain and the analyte into a change in the fluorescence lifetime of the chromophore.

### Construction and characterization of FLIM biosensors for citrate and glucose

To further prove the versatility of our engineering approach for mTQ2-based FLIM biosensors, we developed FLIM biosensors for citrate and glucose using qmTQ2-ATP-0 backbone.

To develop a citrate FLIM biosensor, the sensing domain for citrate (the citrate-binding periplasmic domain (CitAP) of *Klebsiella pneumoniae* CitA) (Zhao et al., 2020) was integrated into mTQ2 between Tyr-145 and Phe-146. The resulting qmTQ2-citrate-0 (quantitative mTurquoise2-based citrate biosensor version 0) showed a 5% increase in fluorescence intensity in the presence of 20 mM citrate. Facile linker optimization was conducted to improve the dynamic range, resulting in a mutant that exhibited a 15% increase in fluorescence intensity and a 0.7 ns fluorescence lifetime change upon exposure to 20 mM citrate. This mutant, named qmTQ2-citrate biosensor, contains a 5-amino-acids (ALKKE) insertion in both linker 1 and linker 2 (Figure 3A). The emission and excitation spectra of qmTQ2-citrate biosensor increased in amplitude by 19% upon the citrate saturation (Figure 3B), with the fluorescence lifetime of qmTQ2-citrate biosensor increasing from 2.2 ns (citrate-free state) to 3.0 ns (citrate-bound state) (Figure 3D). A minimal change in the absorption spectra of qmTQ2-citrate biosensor was observed with or without citrate (Figure 3C). The qmTQ2-citrate biosensor changes its fluorescence lifetime in response to citrate in a dose-dependent manner (Figure 3E), with the estimated K_d_ value being 4.6 ± 0.29 mM (Figure S5). Notably, the fluorescence lifetime of qmTQ2-citrate biosensor remains insensitive to pH between 7.0 and 7.5 (Figure 3F). Moreover, qmTQ2-citrate biosensor exhibited specificity to citrate over other metabolites (Figure 3G).

**Figure 3.**
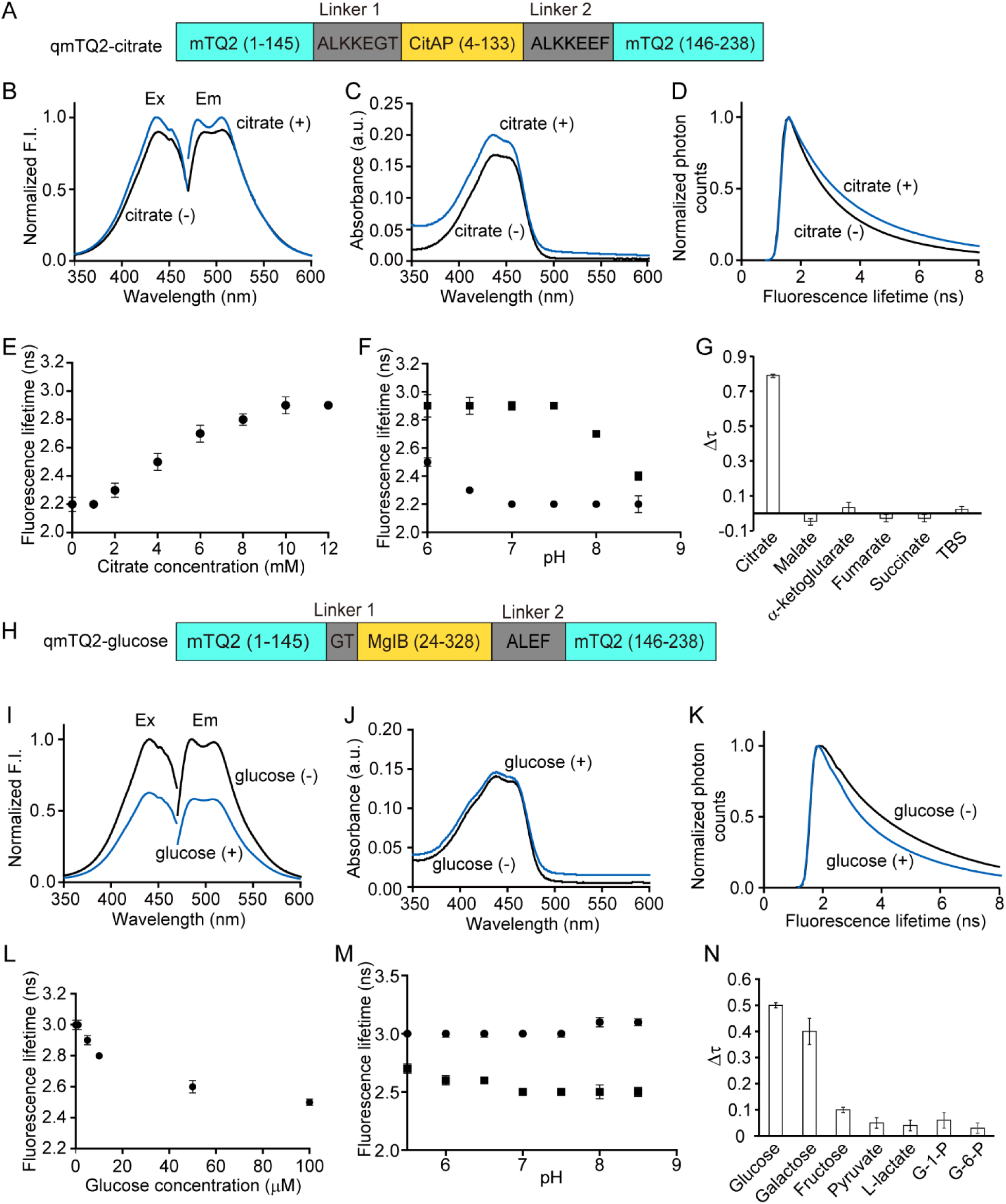
Construction and characterization of qmTQ2-citrate and qmTQ2-glucose biosensors using mTQ2 platform. (A) Schematic domain structure of the qmTQ2-citrate biosensor. (B–D) The emission and excitation spectra (B), the absorption spectra (C) and the fluorescence decay curve (D) of qmTQ2-citrate biosensor in the presence (blue) and absence (black) of 20 mM citrate. (E) The dose-response curve of the qmTQ2-citrate biosensor to citrate in solution. The data represents means ± SD (n = 5). (F) Effect of pH on the fluorescence lifetime of qmTQ2-citrate biosensor in the presence and absence of 20 mM citrate. The data represents means ± SD (n = 5). (G) Specificity of qmTQ2-citrate biosensor to citrate and other metabolites. Δτ represents the dynamic range obtained with the presence and absence of 20 mM metabolites in TBS buffer. The data represents means ± SD (n = 5). (H) Schematic domain structure of qmTQ2-glucose biosensor. (I–K) The emission and excitation spectra (I), the absorption spectra (J) and the fluorescence decay curve (K) of qmTQ2-glucose biosensor in the presence (blue) and absence (black) of 10 mM glucose. (L) Dose-responsive curve of qmTQ2-glucose biosensor to glucose in solution. The data represents means ± SD (n = 5). (M) Effect of pH on the fluorescence lifetime of qmTQ2-glucose biosensor in the presence and absence of 10 mM glucose. The data represents means ± SD (n = 5). (N) Specificity of qmTQ2-glucose biosensor to monosaccharides and glucose metabolism-related molecules. Δτ represents the dynamic range obtained with the presence and absence of 150 μM these molecules. The data represents means ± SD (n = 5).

The epsilon subunit of qmTQ2-ATP-0 backbone was replaced by the glucose sensing domain, bacterial D-galactose-binding periplasmic protein (MglB) (Mita et al., 2022), for constructing the glucose FLIM biosensor. Subsequent linker screening performed on the resulting qmTQ2-glucose-0 (quantitative mTurquoise2-based glucose biosensor version 0) yielded a mutant with a 16% decrease in fluorescence intensity and a 0.3 ns fluorescence lifetime change in response to 10 mM glucose. This mutant, featuring a 2-amino-acid (AL) insertion in linker 2, was designated as qmTQ2-glucose biosensor (Figure 3H). Upon the addition of 10 mM glucose, the emission and excitation spectra of qmTQ2-glucose biosensor decreased in amplitude by 40% (Figure 3I) accompanied by a fluorescence lifetime change from 3.0 ns (glucose-free state) to 2.5 ns (glucose-bound state) (Figure 3K). However, the change in the absorption spectra of the qmTQ2-glucose biosensor was minimal with or without glucose (Figure 3J). qmTQ2-glucose biosensor changes its fluorescence lifetime in response to glucose in a dose-dependent manner (Figure 3L), and the K_d_ value was estimated to be 13.8 ± 2 μM (Figure S6). Additionally, the fluorescence lifetime of qmTQ2-glucose biosensor remains insensitive to pH between 7.0 and 7.5 (Figure 3M). Furthermore, qmTQ2-glucose biosensor showed specificity to glucose and galactose over other monosaccharides and glucose metabolism-related molecules (Figure 3N).

Notably, only facile linker screening was performed in developing qmTQ2-citrate and qmTQ2-glucose, further demonstrating that qmTQ2-ATP-0 backbone is an excellent platform for engineering FLIM biosensors. By replacing the epsilon subunit with analyte sensing domains, combined with linker screening and site-saturation mutagenesis targeting Tyr-145 and linker 1, mTQ2-based FLIM biosensors for various analytes could be developed.

### Dual-color imaging with qmTQ2-cAMP and Rcamp1h biosensors

A single-FP FLIM biosensor has a significant advantage over a FRET-FLIM biosensor in multicolor imaging, requiring only a single wavelength for detection. To explore the potential of dual-color imaging, we prepared HeLa cells that co-expressed qmTQ2-cAMP biosensor along with a red calcium FLIM biosensor, RCaMP1h (Akerboom et al., 2013). Upon simultaneous treatment of the HeLa cells with 100 μM isoproterenol and 50 μM histamine dihydrochloride, the fluorescence lifetime of both qmTQ2-cAMP and RCaMP1h increased (Figures 4A–4C). The increase in RCaMP1h lifetime indicated an elevated Ca^2+^ concentration following histamine stimulation, while the elevation in qmTQ2-cAMP lifetime reflected an augmented cAMP level induced by isoproterenol treatment. The results revealed the feasibility of dual-color imaging using the qmTQ2-cAMP biosensor alongside other biosensors.

**Figure 4.**
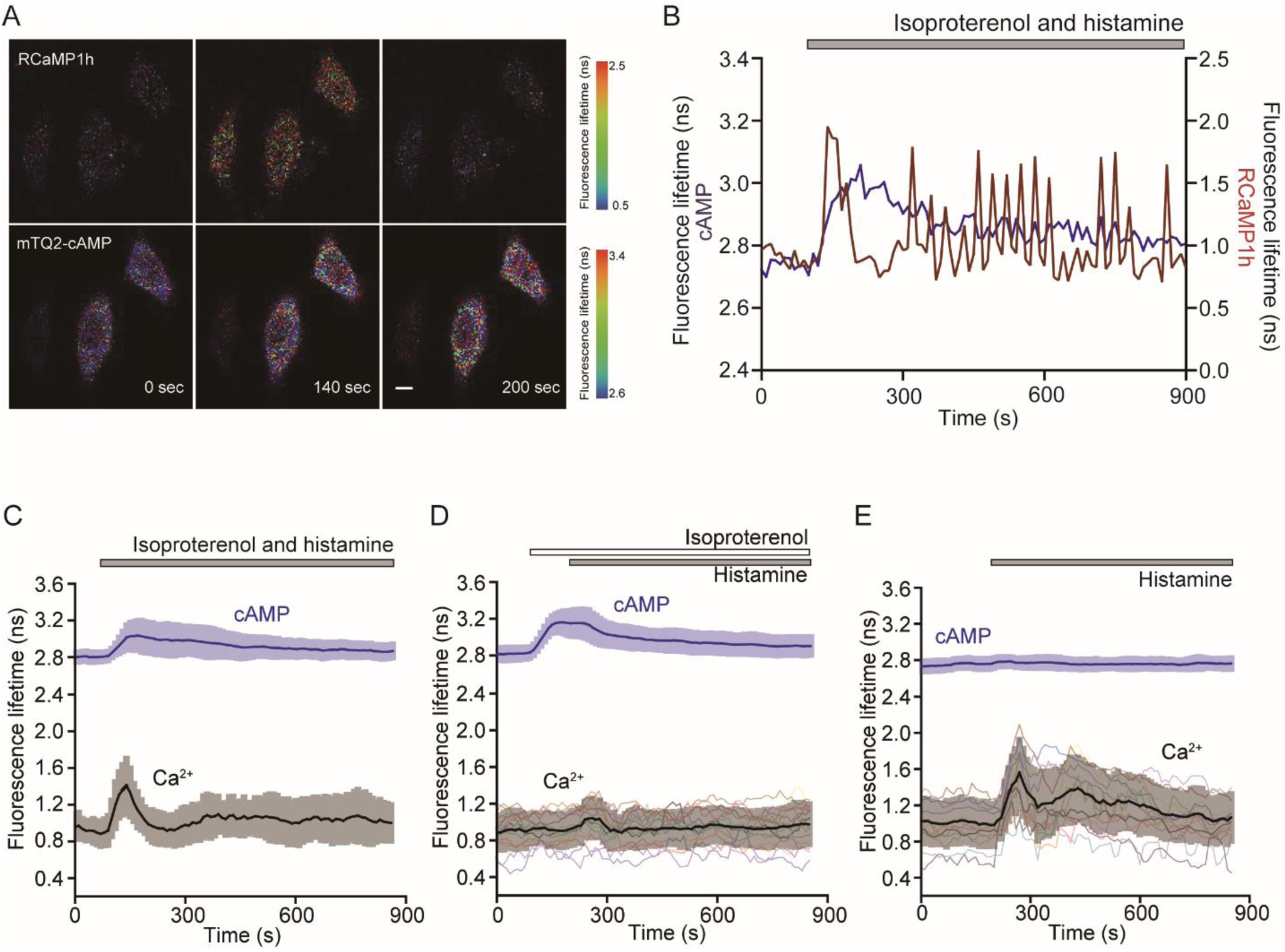
Dual color imaging of HeLa cells expressing the qmTQ2-cAMP and RCaMP1h biosensors. (A) Sequential pseudo-color images of HeLa cells co-expressing qmTQ2-cAMP and RCaMP1h biosensors in response to simultaneous 50 µM histamine dihydrochloride and 100 µM isoproterenol treatment. (B–C) Typical time course (B) and average trace (C) of fluorescence lifetime changes of qmTQ2-cAMP and RCaMP1h biosensors in response to simultaneous histamine and isoproterenol stimulation (n = 12). (D–E) Inhibition of histamine induced calcium oscillations by isoproterenol. Time course of HeLa cells stimulated with 10 μM histamine solely (C) or after pre-treatment with 100 μM isoproterenol (D). Thick lines represent the average traces with qmTQ2-cAMP biosensor shown in purple and the RCaMP1h biosensor in gray. and colorful thin lines represent individual traces (n = 15), scale bar:10 μm.

Histamine induces calcium oscillation via H1 receptor binding, leading to the generation of inositol 1,4,5-triphosphate (IP3). IP3 then binds to IP3 receptors on the endoplasmic reticulum (ER) membrane, prompting the release of calcium ions from the ER lumen into the cytoplasm. Our results suggested that the surge in cAMP levels might inhibit the Ca^2+^ signals triggered by histamine, as evidenced by the simultaneous increase in qmTQ2-cAMP lifetime and a decrease in RCaMP1h lifetime (Figure S8). Previous studies have shown that isoproterenol suppresses histamine-evoked Ca2+ signals in bronchial airway smooth muscle cells through the cAMP and protein kinase A (PKA) pathway (Dale et al., 2018). To further validate the Inhibitory effect of isoproterenol on histamine-induced calcium oscillation in HeLa cells, we performed sequential treatments on HeLa cells co-expressing qmTQ2-cAMP and RCaMP1h biosensors with 100 μM isoproterenol followed by 10 μM histamine. In cells treated solely with histamine, the fluorescence lifetime of RCaMP1h at its maximal peak was 1.5 ± 0.3 ns. However, pre-treatment with isoproterenol significantly reduced this peak lifetime to around 1.04 ± 0.2 ns (Figures 4D and 4E). This result suggested that isoproterenol inhibited histamine-evoked Ca^2+^ signals by reducing its maximal peak rise in HeLa cells. Given the known crosstalk between Ca^2+^ and cAMP signaling pathways at multiple levels (Hofer, 2012), simultaneously monitoring both molecules will offer deeper insights into the intricate crosstalk between these two messengers.

## Discussion

In this paper, by adopting the conventional fluorescence intensity-based biosensor screening strategy, we successfully developed mTQ2-based FLIM biosensors for ATP, cAMP, glucose and citrate. We validated qmTQ2-ATP and qmTQ2-cAMP biosensors in live cell applications. Additionally, we conducted dual-color imaging using qmTQ2-cAMP and Rcamp1h, demonstrating their efficacy. Notably, our results indicated that isoproterenol attenuated histamine-induced Ca^2+^ signals in HeLa cells by reducing their maximum peak rise. The interplay between Ca^2+^ and cAMP signaling is a commonly discussed topic in various research fields, including neurobiology and physiology (Yokoyama et al., 2024). The quantitative imaging method presented in this study will significantly contribute to elucidating the reciprocal regulation between Ca^2+^ and cAMP signaling circuits. Recent advancements in multiplexing, especially with colorful intensiometric biosensors, facilitate the simultaneous observation of the spatiotemporal dynamics of signaling molecules within the same cell (Arai et al., 2018; Zhao et al., 2011). Despite the expanded availability of biosensors, their findings have only captured relative changes in signaling molecules over time. Conversely, multiplex FLIM imaging enables the concurrent quantification of various cellular signaling molecules, providing reliable concentration information. In addition, mTQ2-based biosensors would be suitable for in vivo imaging using two-photon excitation. Our mTQ2-based FLIM biosensors could offer quantitative insights both at the single-cell level and in vivo.

We further addressed the underlying sensing mechanism of mTQ2-based FLIM biosensors. Using fluorescence lifetime and quantum yield, we estimated the radiative rate constant (*k*_r_) and the non-radiative rate constant (*k*_nr_) of qmTQ2-ATP biosensor, along with two mutants qmTQ2-ATP-0.2 and qmTQ2-ATP-0.3 (Table S2). Following ATP binding, the *k*_nr_ of the qmTQ2-ATP mutants increased significantly, while minimal changes were observed in the *k*_r_, indicating that ATP-induced conformation change would increase the non-radiative decay with little effects on the radiative pathway. Consequently, the fluorescence lifetime of the qmTQ2-ATP mutants decreases upon ATP binding, as fluorescence lifetime is inversely proportional to the sum of *k*_nr_ and *k*_r_ of the chromophore. Notably, similar trends were observed with other mTQ2-based biosensors for detecting cAMP, glucose, and citrate (Table 1). In these instances, alterations in fluorescence lifetime induced by analyte binding were primarily driven by changes in *k*_nr_, while *k*_r_ and the molar absorption coefficient were barely affected.

To delve deeper into the mechanism behind the fluorescence lifetime responsiveness of mTQ2-based FLIM biosensors, understanding the structure of mTQ2 is crucial. The long mono-exponential fluorescence lifetime of mTQ2 was achieved by ensuring the planarity of the chromophore and restricting its rotational freedom through strong van der Waals interactions with surrounding amino acids (Goedhart et al., 2012). Moreover, mTQ2 has an exceptionally low pKa of 3.1, so the protonation state of its chromophore is unlikely to be influenced by the analyte-induced conformational changes in the biosensor (van der Linden et al., 2021). Consequently, the changes in the molar extinction coefficient (ε) would be minimal upon analyte binding, as supported by our experimental data (Tables 1 and S2). Taking these features into account, a key aspect of the sensing mechanism in mTQ2-based turn-off FLIM biosensors, such as qmTQ2-ATP and qmTQ2-glucose, could be attributed to the increased mobility of the chromophore upon analyte binding. Specifically, the accelerated free rotation of the chromophore, without concurrent alteration in its charged states, facilitates the dissipation of energy released from the excited state through a vibration relaxation process. Conversely, for mTQ2-based turn-on biosensors like qmTQ2-cAMP and qmTQ2-citrate, the sensing mechanism involves a reduction in chromophore mobility upon analyte binding. These photochemical attributes of mTQ2-based FLIM biosensors, wherein fluorescence intensity changes occur while the ε value (absorption) remains unaffected, could serve as a distinguishing feature for identifying FLIM-based biosensors. Furthermore, it is worth mentioning that mTQ2, with its long fluorescence lifetime, might have the potential for developing FLIM biosensors with expanded dynamic ranges. This is ascribed to the unique molecular environment surrounding the chromophore, offering a versatile platform for the development of various FLIM biosensors.

It should also be noted here that the biosensors developed by this strategy have dual functionality. They respond to the analytes by both fluorescence intensity and lifetime. FLIM set-up is not required to use these biosensors. The same biosensor can be used with conventional fluorescence microscopes for easier intensiometric experiments. Researchers can then use the FLIM setup for more precise analyses.

These observations suggest that other FPs with high quantum yield would also serve as good scaffolds for designing FLIM-based biosensors. Recently reported bright and photostable YFP variant StayGold and its derivatives (Ando et al., 2024; Hirano et al., 2022; Zhang et al., 2024), with a reported quantum yield of 0.93, show promise in this regard. StayGold, with its spectral separation from mTQ2, is well-suited for simultaneous measurements, and its further development is highly anticipated.

Overall, mTQ2-based FLIM biosensors provide novel tools for visualizing and quantitatively assessing intracellular dynamics of ATP, cAMP, citrate, and glucose. Additionally, we demonstrated that the qmTQ2-ATP-0 backbone is a universal platform for developing mTQ2-based biosensors. Moreover, we proposed a facile screening strategy for constructing fluorescence lifetime-based biosensors using microplate fluorescence readers, which is relatively low-cost and widely accessible. Our research will significantly facilitate the development of FLIM biosensors for quantitative fluorescence imaging in the future.

## Acknowledgments

We thank Dr. Takashi Jin (RIKEN) for his generous support in measuring the absorption spectrum and quantum yield. We thank Dr. Kazunari Mouri (National Institute of Information and Communications Technology, Kobe frontier research center) and Dr. Cong Vu (Kanazawa University) for useful suggestions. We also thank our lab members for the discussion, Ms. Xu Shang Dan, Yukiko Onishi, Mikako Hayashi, Sayuri Yamamoto, and Jyunko Asada for technical support, and Ms. Manaho Kakiuchi and Tomko Furuya for secretarial assistance. C.Z. was supported by JST SPRING program (JPMJSP2135). This work was also supported by JSPS through KAKENHI grants (19H03394, 19H05794, 19H05795, 22H02798, and 22H04926) to Y.O. and World Premier International Research Center Initiative (WPI) to S.A., by JST through FOREST Program (JPMJFR201E) to S.A. CREST (JPMJCR1852 and JPMJCR20E2) to Y.O., and Moonshot R&D grant (JPMJMS2025-14) to Y.O.

## Author contributions

Y.O. and S.A. conceived the project. C.Z. designed and conducted the experiments, analyzed the data, and wrote the manuscript. All the authors reviewed the manuscript.

