## Supplementary information for "A versatile platform for single fluorescent protein-based fluorescence lifetime biosensors"

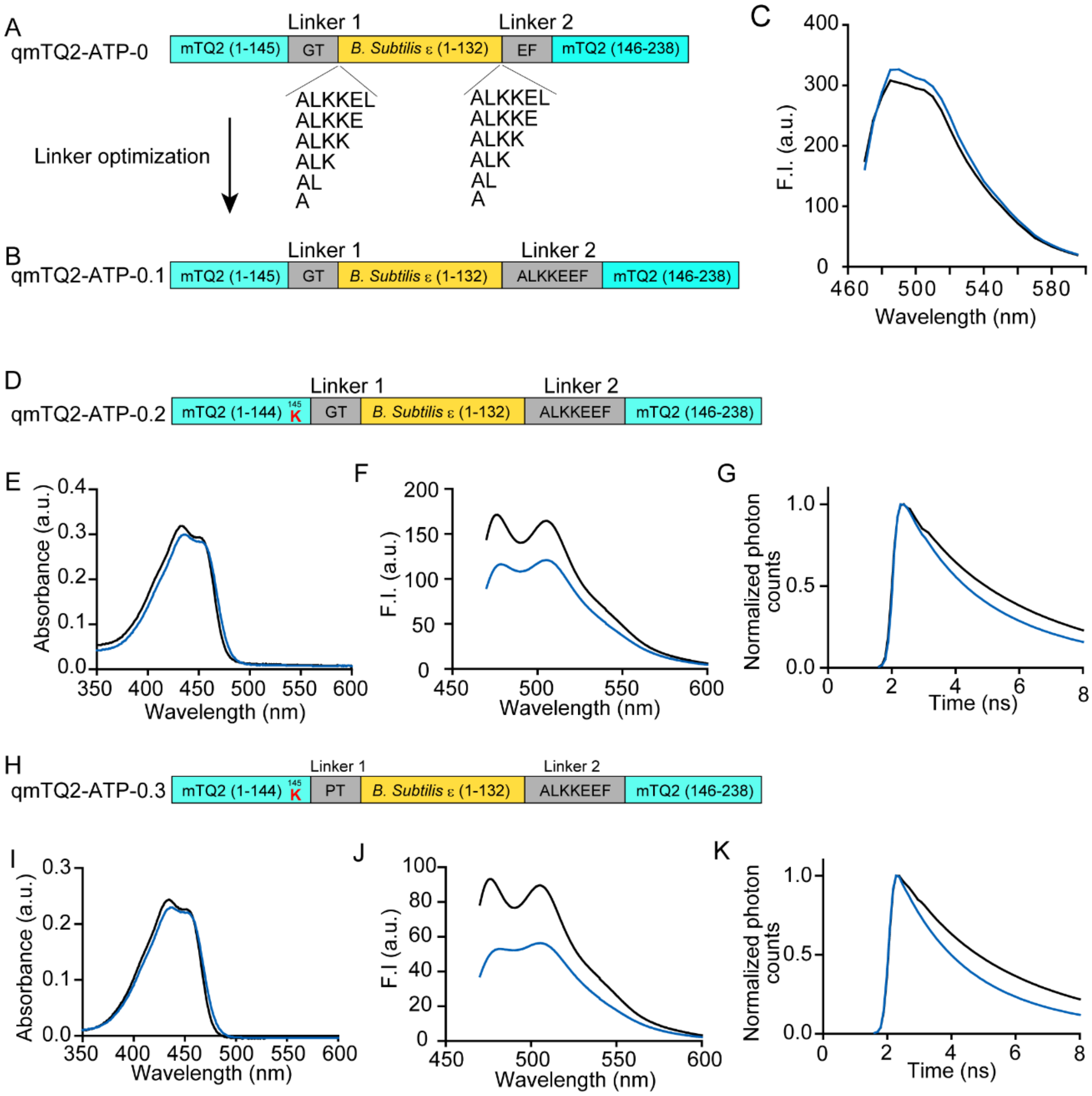


**Supplementary Figure 1**. Illustrations of the detailed screening process for the qmTQ2-ATP sensor. (A–C) Schematic drawings of the linker optimization process. The domain structures for qmTQ2-ATP-0 (A) and qmTQ2-ATP-0.1 (B). (C) The emission spectra of qmTQ2-ATP-0.1 with (blue line) and without (black line) 10 mM ATP. (D) Domain structure illustration for qmTQ2-ATP-0.2. Absorption spectra (E), emission spectra (F) and fluorescence decay curve (G) of qmTQ2-ATP-0.2 with (blue line) and without (black line) 10 mM ATP. (H) Domain structure illustration for qmTQ2-ATP-0.3. Absorption spectra (I), Emission spectra (J) and fluorescence decay curve (K) of qmTQ2-ATP-0.3 with (blue line) and without (black line) 10 mM ATP. The fluorescence intensity change (ΔF/F) and fluorescence lifetime change (Δτ) of the qmTQ2-ATP mutants were summarized in Supplementary Table 1.

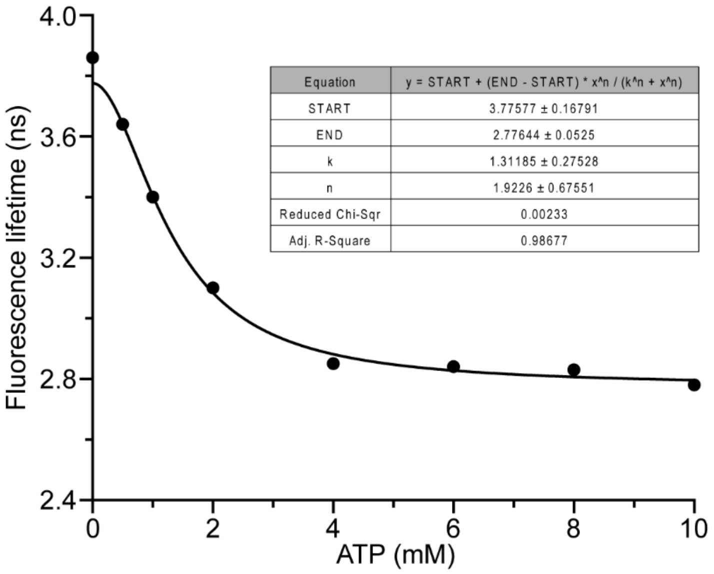


**Supplementary Figure 2.** Curve fitting for the qmTQ2-ATP sensor dose-responsive curve. Curve fitting was performed using modified Hill function with offset (Hill1 mode) in Origin software (OriginPro, Version 2022b. OriginLab Corporation, Northampton, MA, USA). The Hill equation is

$$y = START + (END - START)\left(\frac{x^n}{x^n + k^n}\right)$$

where y is fluorescence lifetime, x is ATP concentration, k is dissociation constant, and n is the Hill coefficient.

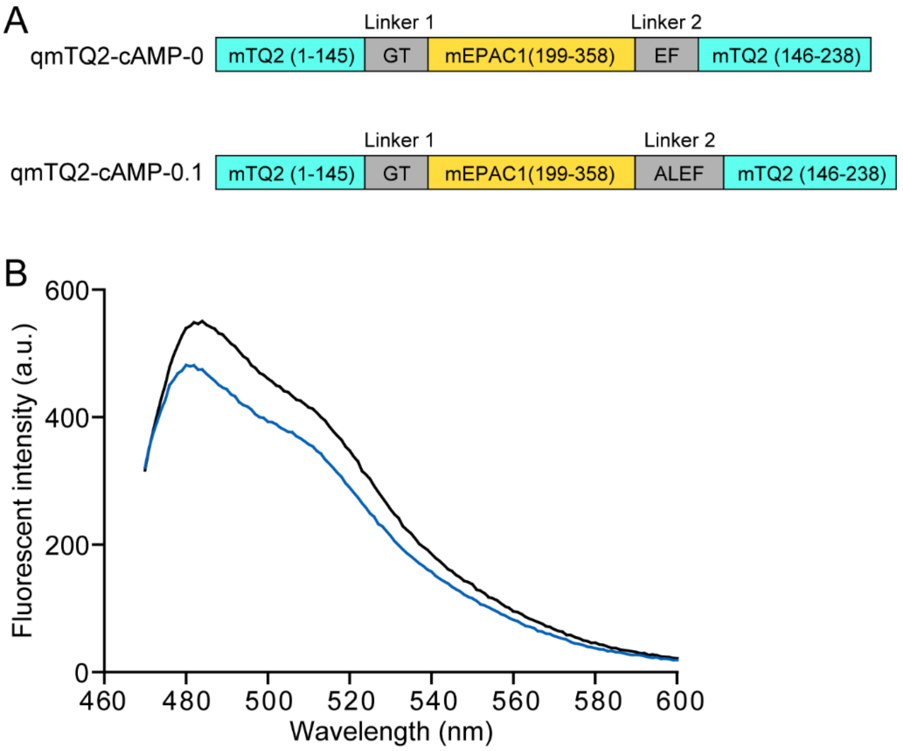


**Supplementary Figure 3**. Linker optimization of the qmTQ2-cAMP sensor. (A) Schematic illustration of the domain structures of the qmTQ2-cAMP-0 and qmTQ2-cAMP-0.1 mutants. (B) The emission spectra of qmTQ2-cAMP-0.1 mutant with (blue line) or without (black line) 1 mM cAMP.

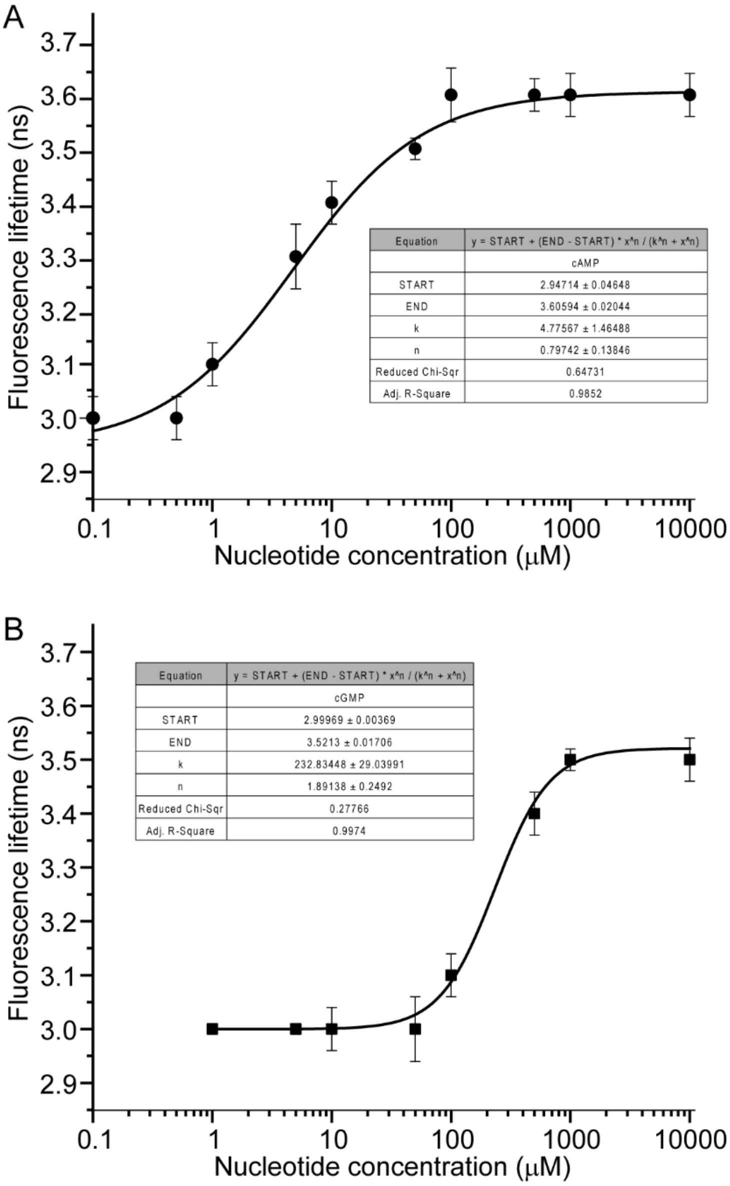


**Supplementary Figure 4.** Curve fitting of the dose-responsive curves of the qmTQ2-cAMP sensor to cAMP (A) and cGMP (B). The fitting curve were obtained using the same equation as that in supplementary Figure 2.

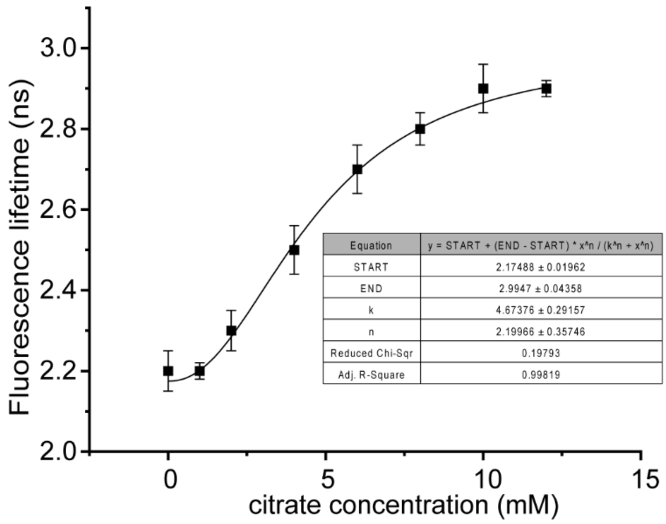


**Supplementary Figure 5.** Curve fitting of the dose-responsive curve of the qmTQ2-citrate sensor. The fitting curve was obtained using the same equation as that in Supplementary Figure 2.

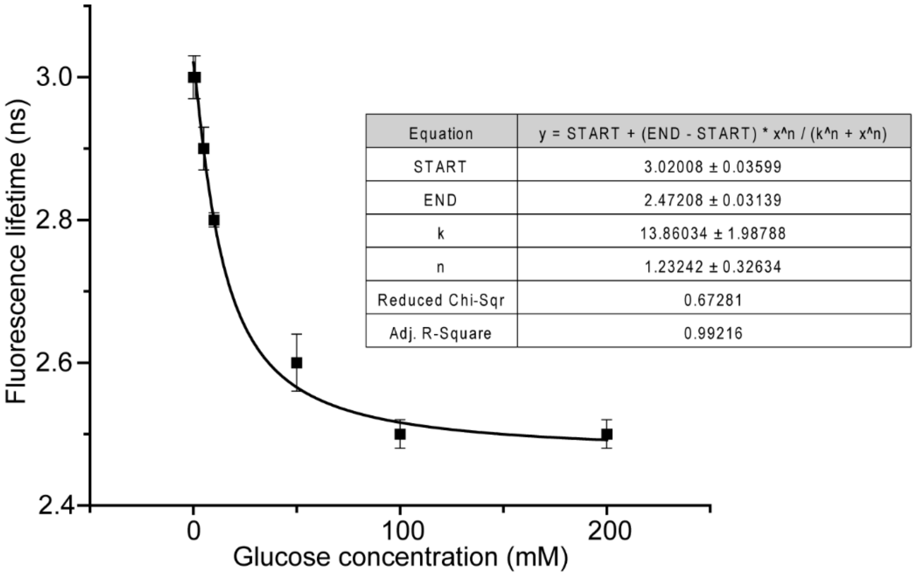


**Supplementary Figure 6.** Curve fitting of the dose-responsive curve of the qmTQ2-glucose sensor. The fitting curve was obtained using the same equation as that in Supplementary Figure 2.

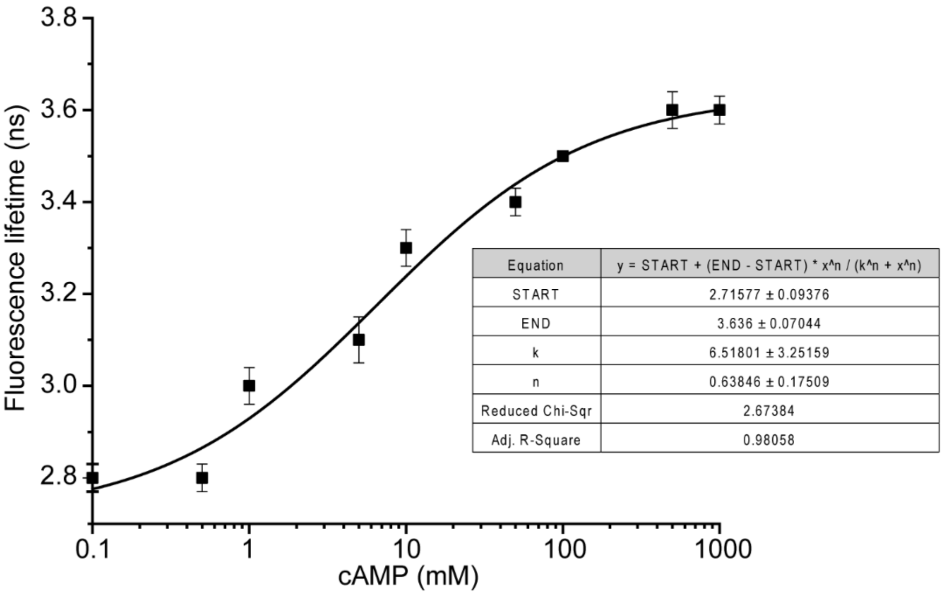


**Supplementary Figure 7.** Calibration curve and curve fitting of the qmTQ2-cAMP sensor in permeabilized HeLa cells. The fitting curve was obtained using the same equation as that in Supplementary Figure 2.

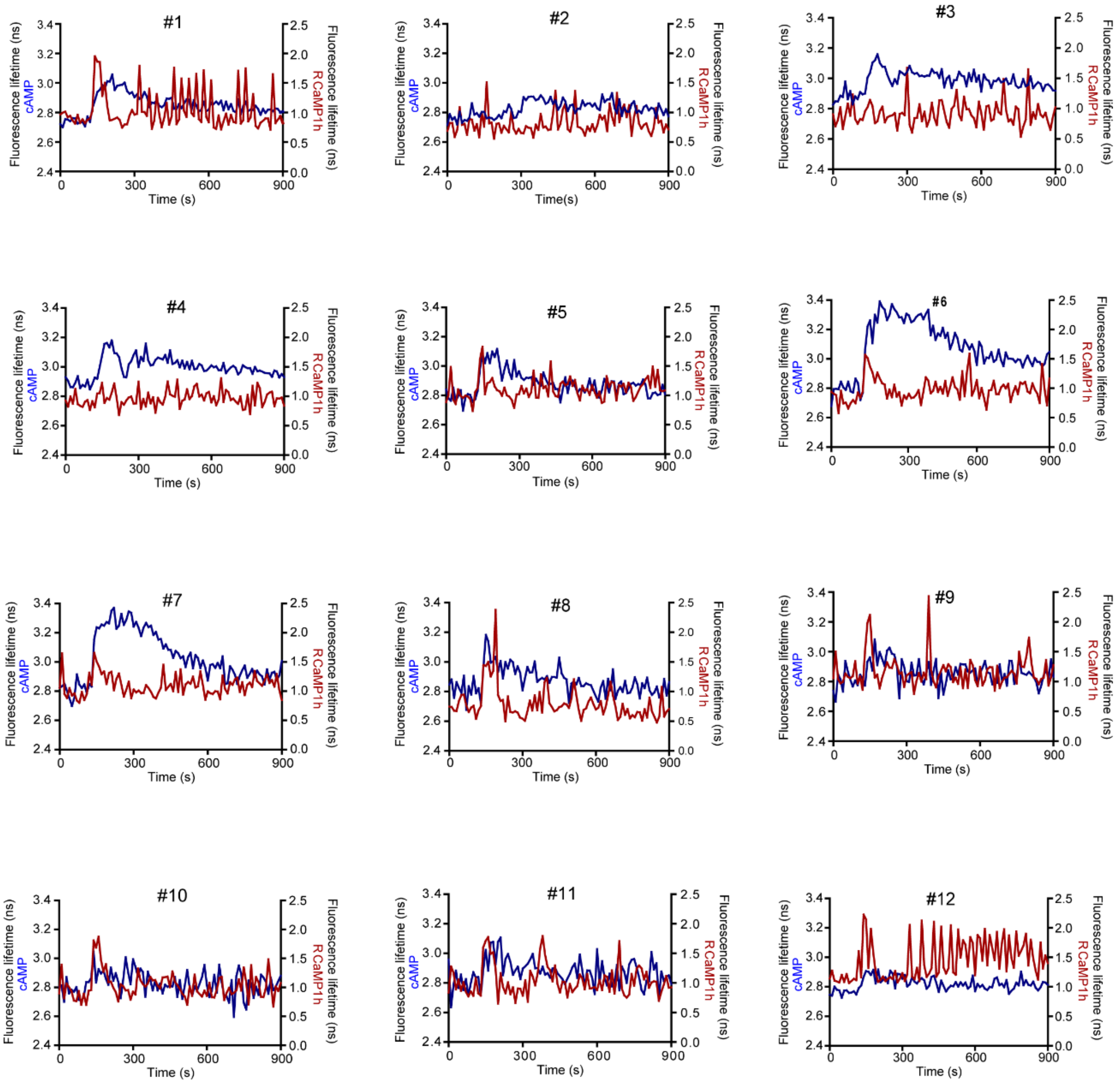


**Supplementary Figure 8**. Summary for all fluorescence lifetime changes traces for the qmTQ2-cAMP and Rcamp1h sensors obtained under the experimental condition for Fig.4 A–C. HeLa cells were simultaneously stimulated with 50 μM histamine and 100 μM isoproterenol 120 seconds after the start of acquisition.

**Supplementary Table 1: Dynamic ranges of qmTQ2-ATP mutants**

| **Tyr-145** | **Linker 1** | **$\Delta F/F_0$** | **$\Delta\tau$ (ns)** |
|---|---|---|---|
| W | RG | 0.19 | 0.3 |
| R | GT | 0.21 | 0.5 |
| K | GT | 0.26 | 0.6 |
| K | PT | 0.34 | 0.9 |
| K | RT | 0.31 | 0.7 |
| K | KT | 0.33 | 0.8 |
| K | PV | 0.4 | 1.1 |
| K | PE | 0.4 | 1.0 |

Fluorescence intensity dynamic range ($\Delta F/F_0$) and fluorescence lifetime change ($\Delta\tau$) of qmTQ2-ATP mutants were obtained with purified protein. Amino acids at Tyr-145 and linker 1 of qmTQ2-ATP mutants are indicated.

**Supplementary table 2. Photophysical characterization of qmTQ2-ATP mutants**

| Name | 10 mM ATP | $\tau$ (ns) | $\varepsilon$ ($\times10^4$ $M^{-1}$ $cm^{-1}$) | QY | $K_r$ (x$10^8$; $s^{-1}$) | $K_{nr}$ (x$10^8$; $s^{-1}$) |
|---|---|---|---|---|---|---|
| qmTQ2-ATP-0.2 | + | 3.2 | 2.54 | 0.54 | 1.65 | 1.41 |
| | - | 3.8 | 2.61 | 0.70 | 1.79 | 0.78 |
| qmTQ2-ATP-0.3 | + | 2.9 | 2.56 | 0.46 | 1.56 | 1.86 |
| | - | 3.8 | 2.64 | 0.66 | 1.73 | 0.88 |
| qmTQ2-ATP | + | 2.7 | 2.61 | 0.47 | 1.69 | 1.90 |
| | - | 3.8 | 2.69 | 0.71 | 1.83 | 0.76 |

$\tau$, Fluorescence lifetime; $\varepsilon$, Molar absorption coefficient; QY, Quantum yield.
